# Continuous *in vivo* metabolism by NMR

**DOI:** 10.1101/501577

**Authors:** Michael T. Judge, Yue Wu, Fariba Tayyari, Ayuna Hattori, John Glushka, Takahiro Ito, Jonathan Arnold, Arthur S. Edison

## Abstract

Dense time-series metabolomics data are essential for unraveling the underlying dynamic properties of metabolism. Here we extend high-resolution-magic angle spinning (HR-MAS) to enable continuous *in vivo* monitoring of metabolism by NMR (CIVM-NMR) and provide analysis tools for these data. First, we reproduced a result in human chronic lymphoid leukemia cells by using isotope-edited CIVM-NMR to rapidly and unambiguously demonstrate unidirectional flux in branched-chain amino acid metabolism. We then collected untargeted CIVM-NMR datasets for *Neurospora crassa*, a classic multicellular model organism, and uncovered dynamics between central carbon metabolism, amino acid metabolism, energy storage molecules, and lipid and cell wall precursors. Virtually no sample preparation was required to yield a dynamic metabolic fingerprint over hours to days at ^~^4-min temporal resolution with little noise. CIVM-NMR is simple and readily adapted to different types of cells and microorganisms, offering an experimental complement to kinetic models of metabolism for diverse biological systems.

---

Metabolic time-series data are invaluable for the development and validation of high-quality models that accurately describe the dynamics of metabolism^1-3^. Information about the metabolic state of an organism typically requires extensive time, resources, and sample material. As such, researchers must choose between variables such as the number of replicates, the number of time points, and the time resolution for time-series. Furthermore, traditional metabolomics experimental designs face the challenges of extraction biases^4^ and the confounding of biological and analytical variance^5^. While many studies employ sample preparation and extraction approaches effectively, direct or *in vivo* measurements are fundamentally simpler to obtain and interpret. Likewise, while carefully designed^6^ and executed studies with large sample sizes undeniably yield powerful insights into the dynamics of biological systems^7-9^, continuous and repeated measurements on the same living sample are invaluable for monitoring and confirming these dynamics.

Small molecules and their fluxes have been measured *in vivo* using NMR^10^, and methods have recently been developed that begin to address the need for a continuous time dimension in metabolomics data. For example, long-standing flow NMR techniques allow monitoring of secretion and uptake of extracellular metabolites for organisms grown in liquid culture^10^. Link et al. recently achieved high temporal resolution on many metabolites by developing an automated real-time metabolomics platform that samples liquid cultures of single cells and directly injects them onto a time-of-flight mass spectrometer every 15-30s^11^. The group have more recently probed the interactions between biomass synthesis and cell division in *E. coli* using this method^12^. Koczula et al. conducted *in vivo* measurements changes in media composition with 4-8 min resolution for chronic lymphoid leukemia. Sedimentation and line broadening are major factors that limit standard NMR measurements of complex samples like cells. Koczula et al. were able to mitigate sedimentation by immobilizing the single cells in agarose^13^.

Alternatively, HR-MAS enables high-resolution NMR measurements on mixed-phase samples such as tissues^14^, or more recently, living organisms^10,15-18^ with minimal line broadening. In this study, we extended HR-MAS to real-time continuous *in vivo* measurements of metabolism in cells. Using isotope editing, CIVM-NMR was able to reproduce and more directly observe a surprising branched-chain amino acid (BCAA) flux result reported last year in human myeloid leukemia cells^19^. We found that CIVM-NMR was not only easier but faster and more conclusive than traditional approaches for flux measurements in human cell cultures. We then applied CIVM-NMR to the multicellular filamentous fungus, *N. crassa*, in both aerobic and anaerobic environments. We observed highly reproducible dynamics in central carbon and amino acid metabolism with ^~^4 min resolution over 11 hours. The continuous nature of these measurements facilitated metabolite annotation, and semi-automated peak tracing provided relative quantification of known and unknown compounds. We developed several new MATLAB functions and workflows, freely available through GitHub, for the analysis and visualization of these novel data. As CIVM-NMR can be applied widely to cells, tissues, and small multicellular organisms, it enables new opportunities in fields such as developmental and chronobiology for monitoring high-resolution metabolic time-series data. Importantly, it will enable more robust and experimentally-based kinetic metabolic models for diverse biological systems.

## RESULTS

### Isotopic CIVM-NMR measurements confirm unidirectional KIV-to-valine flux in ML cells

Branched-chain amino transferase-1 (BCAT1) is a reversible enzyme, but in most cells the reaction degrades BCAAs and makes branched-chain keto acid (BCKA)s. However, we recently demonstrated that (BCKA) transamination by the BCAT1 enzyme builds up the BCAA pool in myeloid leukemia (ML) cells, essentially running in the reverse direction^19^. When α-keto-isovalerate (KIV; one of the substrates of BCAT1) was ^13^C-labeled, valine (the expected product of BCAT1) containing ^13^C accumulated. Labeled KIV was not observed when ^13^C-labeled valine was supplied, indicating a non-canonical, unidirectional flux from KIV to valine^19^. In that study, metabolic fingerprints were acquired via a traditional, labor- and material-intensive sampling scheme involving months of sample preparation and several dozen samples. One reason for the large number of samples in this or similar studies is the biological and technical variation due to sample preparation steps; these factors make it more challenging to compare time-series data without large numbers of replicates. We sought to replicate the result of the original Hattori et al. study using real-time *in vivo* metabolomics.

First, we cultured myeloid leukemia cells as previously described^19^, then pelleted and resuspended them in IMDM media without KIV or valine. Working quickly, we loaded the cells into an HR-MAS rotor and added either ^13^C-labeled KIV or valine to make a total volume of ^~^60 µL, capped the sample, and inserted the rotor into the magnet. We recorded 1D HSQC spectra every 4.2 minutes while spinning at 3500 Hz at the magic angle (54.7°)^14^. A hole in the rotor cap allowed for gas exchange^15^.

By monitoring the intensity of the methyl peaks of both KIV and valine, we observed that ^13^C-labeled KIV decreased in intensity and fell close to the limit of detection within about 60 min (**Fig. 1a**). The ^13^C-labeled valine peak grew with an inversely proportional trajectory, providing real-time, *in vivo* evidence of KIV-to-valine conversion. As the reaction rate depended on the concentration of the cells in the rotor, cell density was adjusted to accommodate measurement of the rapid reaction and provide greater detail about reaction kinetics. As reported previously, labeled KIV was not observed when ^13^C-labeled valine was supplied (**Fig. 1b**). Thus, the original results from Hattori et al. that took months of sample prep and data collection were reproduced with real-time resolution in one afternoon. We did not need to adapt the culture media^11^ or embed the cells^13^ to get these results. Additionally, the combined rate of uptake and conversion of valine could be measured with precision, where measurements at only a few time points were taken previously.

**Figure 1.**
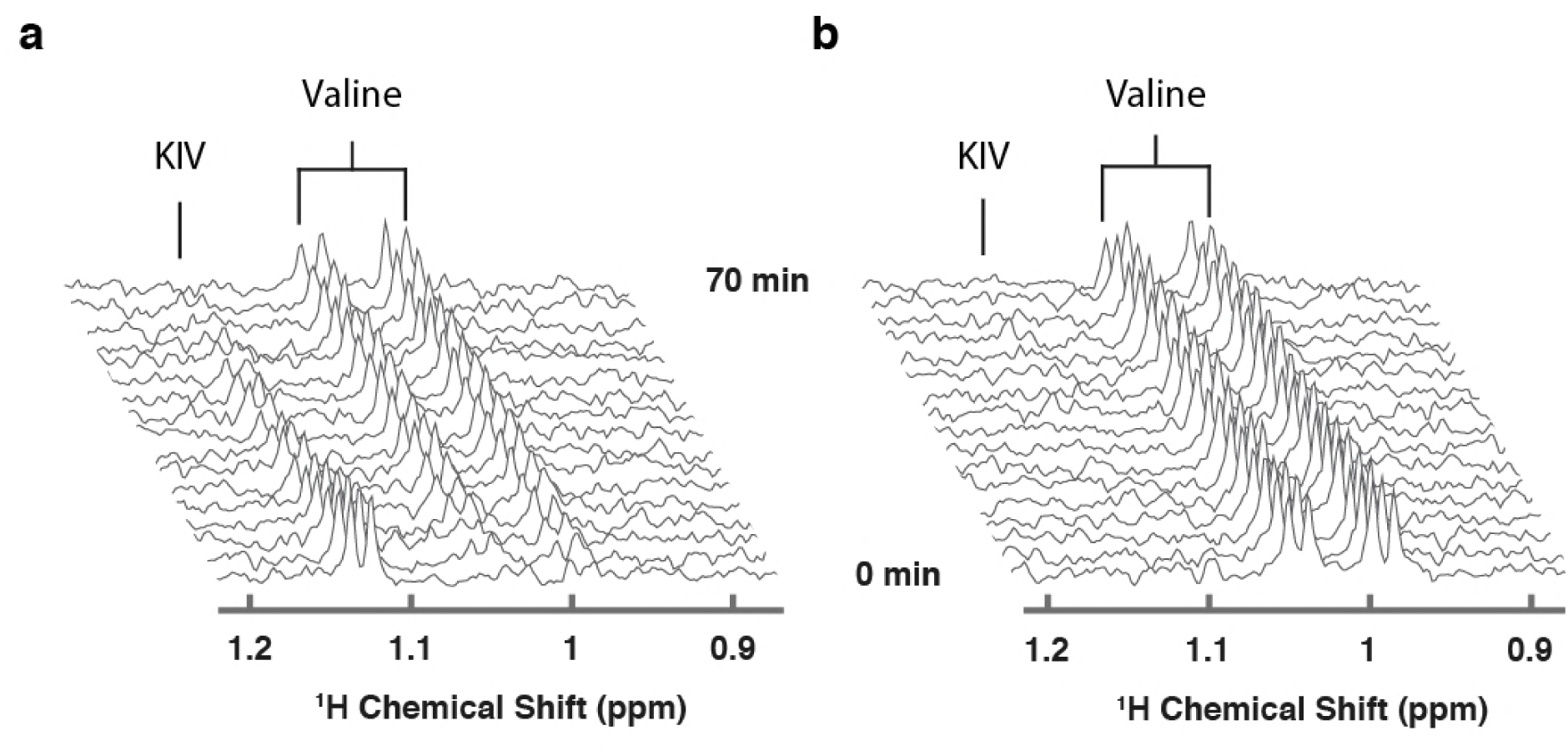
Targeted isotopic CIVM-NMR measurement of metabolic flux in human myeloid leukemia cells. **(a)** ^13^C-labeled keto-isovalerate (KIV) was converted to valine. **(b)** ^13^C-labeled valine was not converted to KIV, confirming unidirectional flux in ML cells.

### Untargeted CIVM-NMR measurements of *N. crassa* growth

Given the utility of CIVM-NMR for the targeted monitoring of known reactions in mammalian cells, we applied it to the continuous measurement of the metabolic dynamics of the filamentous fungus *N. crassa* over 11 h in both aerobic and anaerobic environments. *N. crassa* is an obligate aerobe but will live under low oxygen conditions^20-22^. We grew *N. crassa* tissue in a nutrient-rich liquid medium (**Fig. 2a**). After 30 h, a piece of tissue with a volume of ^~^50 μL was taken from the main mycelial mass, rinsed, and put into a 4-mm HR-MAS rotor with fresh media. The rotor was sealed with a cap with a hole filtered with rayon culture tape punches (“aerobic”)^15^ or no hole (“anaerobic”), placed in the HR-MAS probe, and spun at 6000 Hz at the magic angle for the duration of each experiment (**Fig. 2b**). Each individual scan of a standard 1dnoesypr experiment took ^~^3.97 s. Scans were recorded and summed continuously, and free induction decays (fids) were written to a file once every 64 scans, establishing our shortest temporal resolution at 4.23 min (**Fig. 2c**). After data acquisition, properly phased and Fourier-transformed frequency-domain data were added together to increase the signal-to-noise ratio (S/N) at the expense of temporal resolution (**Fig. 2d**). The organism was assessed for survival after each experiment (ranging between 11 h to 4 days). In every case (n = 9), mycelia did not sediment, were intact, and grew significant hyphae within hours of being placed on standard nutrient agar (**Supplementary Fig. 1**). Thus, *N. crassa* survived the CIVM-NMR experiments and could be used in downstream experiments or processing steps (**Fig. 2e**). Custom shell scripts allowed for batch processing of NMR data (**Fig. 2d**) using NMRPipe^23^. Normalizing to the stable 1 mM DSS reference resonance (0.0 ppm) allowed for relative comparison of peak intensities across time points and samples. To improve S/N, sequential spectra were time-averaged, resulting in 12.7-min temporal resolution for all downstream analyses (**Fig. 2d**).

**Figure 2.**
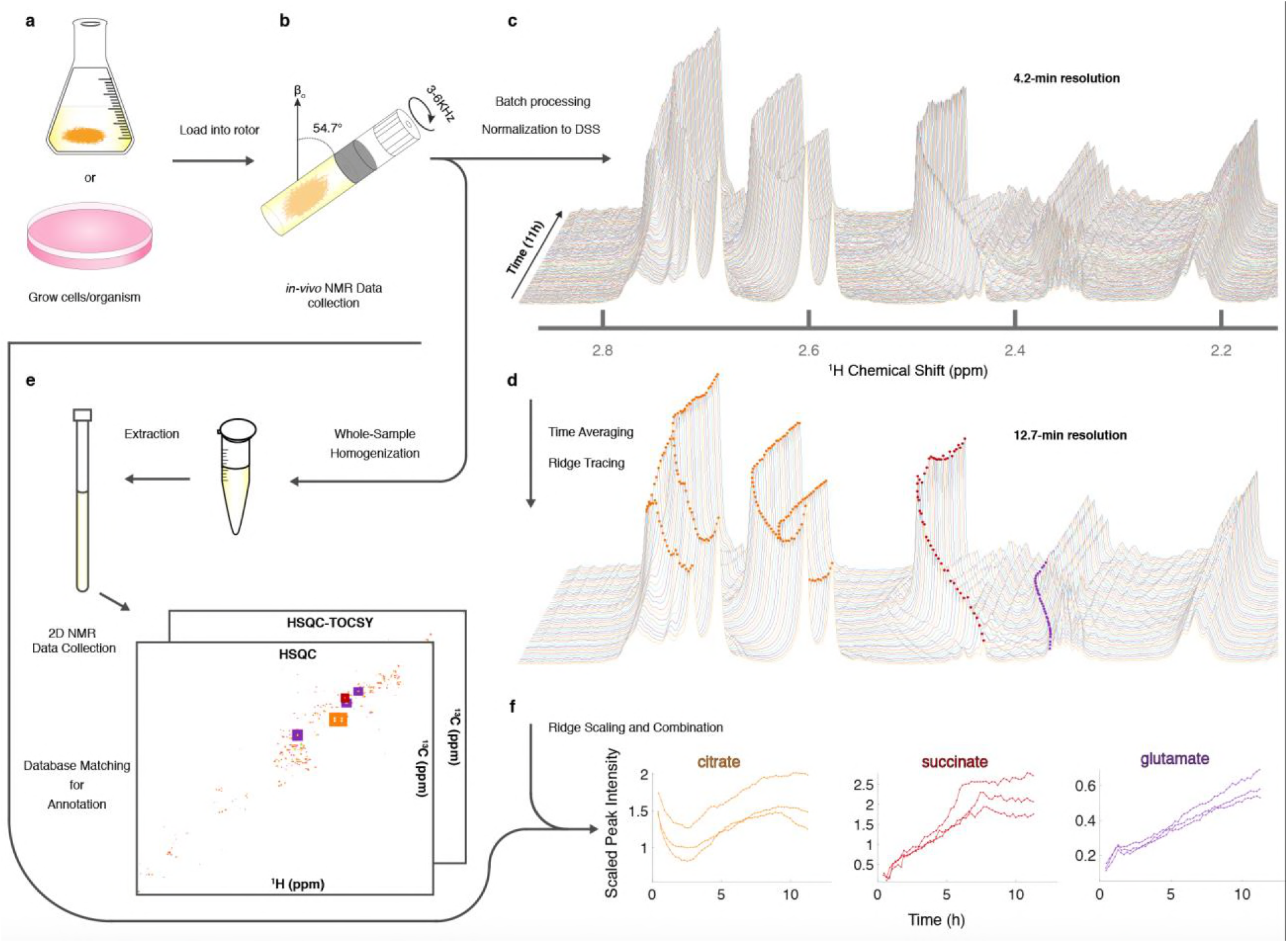
Sample preparation and analysis for CIVM-NMR experiments. (**a**) Samples were first grown to a suitable volume or density in standard media and (**b**) transferred to the HR-MAS rotor (*N. crassa* is shown). Gas composition (e.g. air availability) was altered using a filtered hole or no hole in the cap, and the rotor was spun at the magic angle. NMR data were collected continuously every 4 minutes over the course of hours, then (**c**) processed and normalized to the DSS reference peak (0 ppm) to yield full-resolution data. (**d**) Every three spectra were time-averaged (summed) for improved S/N, and peak intensities were traced across time using ridge tracing to yield relative quantification of metabolites. (**e**) Following HR-MAS, the rotor contents were homogenized, methanol-extracted, and used for 2D NMR analysis for peak annotation by database matching. (**f**) For annotated metabolites with >1 peak (e.g. citrate), the quantified and annotated trajectories (ridges) for each peak were scaled and combined into a single representative trajectory. Trajectories for each annotated compound in 3 aerobic experiments are plotted to compare time series between biological replicates.

To assist with annotation and compound identification, the organism and media were removed at the end of each run, bead-homogenized, and extracted in MeOH (80%) (**Fig. 2e**). Combined supernatants for representative samples were analyzed using ^13^C-HSQC, HSQC-TOCSY, and noesypr1d NMR experiments, and the data were matched to an NMR metabolomics database using COLMARm^24^. Resulting putative identifications were manually assigned confidence scores as described previously^25^. We mapped 34 metabolites with high confidence scores onto the real-time *in vivo* spectra of *N. crassa* (representative annotations, **Fig. 2f**), including multiple amino acids and metabolites involved in the TCA cycle, glycolysis, and fermentation (**Fig. 3a-c; Supplementary Table 1**). Several metabolites overlapped with those found in a previous NMR study in *N. crassa^26^*. We created MATLAB functions for visualization of time series data for samples individually (**Fig. 2c-d**) or as interactive mirror images (**Fig. 3**). We found that the latter approach facilitated comparison between samples, revealing several differences in metabolism between the aerobic and anaerobic conditions (**Fig. 3**) that were reproduced in replicate samples (**Supplementary Fig. 2**).

**Figure 3.**
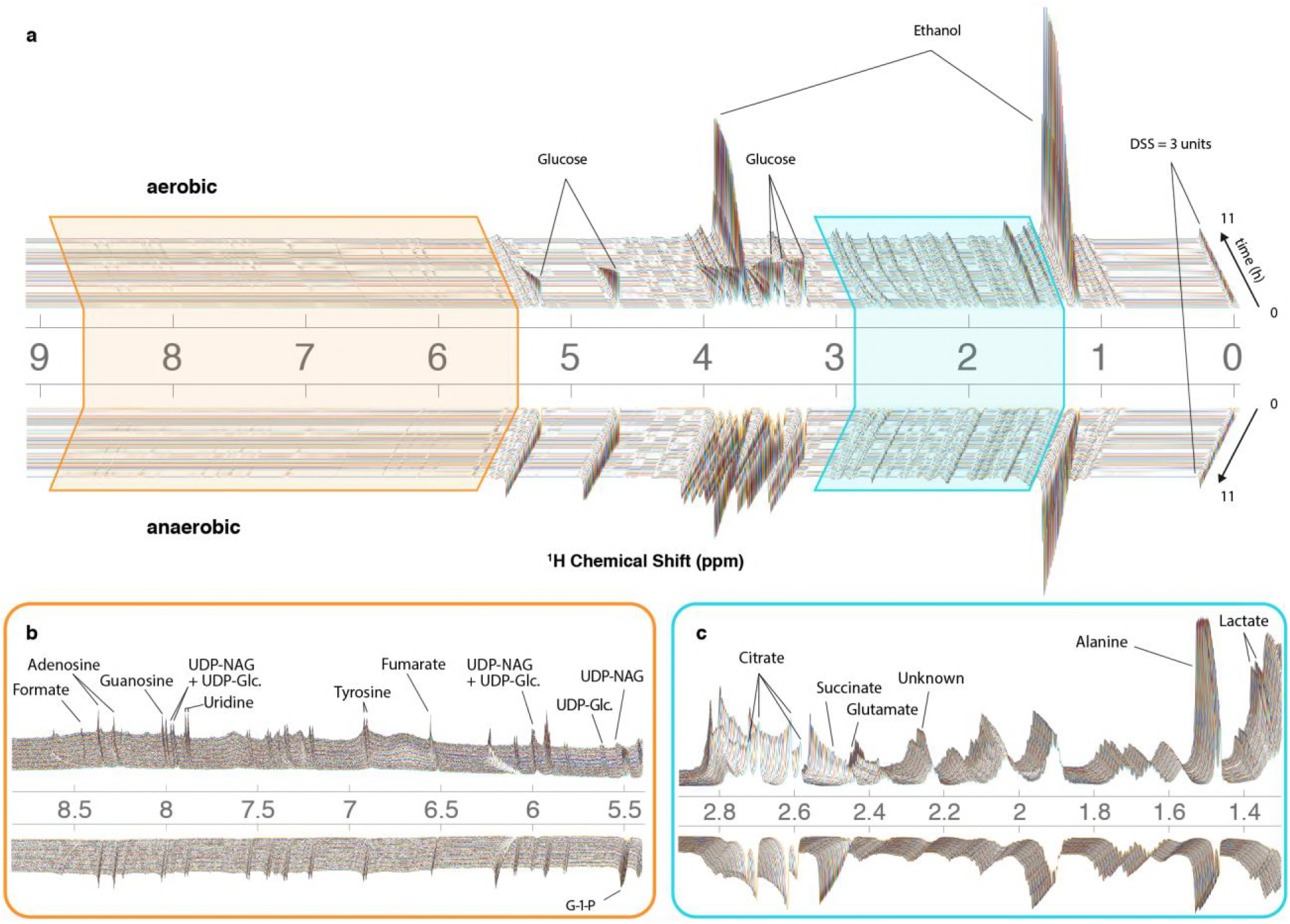
CIVM-NMR measurements of *N. crassa* metabolism under aerobic and anaerobic conditions. ^1^H NMR data for one aerobic replicate (top) and one anaerobic replicate (bottom) plotted interactively as a ‘mirror plot’ for direct comparison between conditions by peak height and position at a given time. To improve the S/N, data were analyzed at 12.7 min resolution. Annotations are shown for select peaks of interest for (**a**) the entire spectrum, and expansions of (**b**) the aromatic region and (**c**) the aliphatic region. Several peaks change position and intensity over the course of the experiments. Abbreviations: UDP-NAG, UDP-N-Acetyl Glucosamine; UDP-Glc., UDP-Glucose; G-1-P, Glucose-1-Phosphate.

### Relative quantification of metabolites by ridge tracing

The 34 compounds that were mapped to *in-vivo* data were assigned a second confidence score for quantifiability (**Online Methods**). For 21 highly scoring metabolites (**Supplementary Table 1**), we obtained relative quantification (**Supplementary Fig. 3**) by tracing peaks across time with a ridge-tracing algorithm (**Figs. 2d, 4a**). With our current algorithm that is limited to peaks with low overlap, we traced over 170 peaks across all of our spectra, including ^~^150 that are currently un-annotated. We combined the information from ridges of sufficient quality when assigned to the same compound (**Fig. 4b**), leveraging the information about compound concentration from multiple measurements. Replicates of dense, continuously repeated measurements on the same sample offer benefits^3^ that would be eliminated by taking time-wise averages or standard errors. We are developing more comprehensive and robust statistical treatments of these unique data within a modeling framework to address this need.

**Figure 4.**
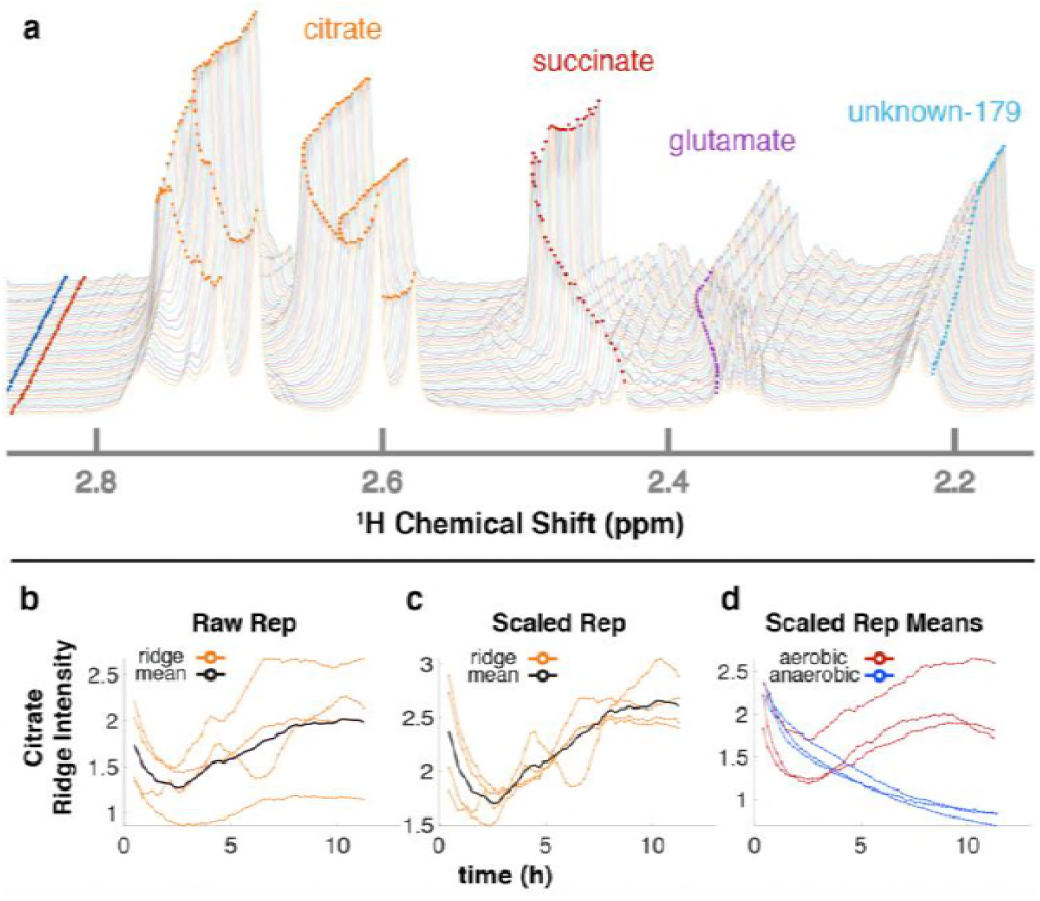
Ridge tracing produces concentration dynamics of metabolites. (**a**) Multiple traced ridges for a single aerobic replicate. Peak maxima at each time point were located using a peak-picking algorithm that includes an adjustable Gaussian filter. Maxima were connected to form ridges along the time dimension using a single linkage hierarchical agglomerative clustering based on Euclidean distances between the points in chemical shift, time, and intensity space. Metabolites typically have several characteristic NMR peaks, e.g. the 4 orange ridges in citrate (**a**). A simple time-wise average represented by the black line in **b** only gives the average intensity over time but loses valuable information on actual dynamic trends. To more accurately extract trends for a particular metabolite, we first integrate each peak in that metabolite over time to obtain its mean value. Then, each peak trajectory is scaled by ratio of the highest mean to its own mean, yielding the 4 orange lines in **c**. The mean of these trajectories is shown in black in **c** and represents the relative concentration over time for that metabolite in that replicate. The 3 aerobic (red) and 3 anaerobic (blue) replicates for citrate are shown in **d**.

### Glucose-dependent changes in pH

NMR chemical shifts are sensitive to pH and metal ion content^27,28^, typically requiring peak alignment algorithms that are prone to creating artifacts. The positions of peaks clearly changed across time in our data (**Figs. 3b-c, 4a**), particularly in the aerobic samples. Because these changes were monitored continuously, peak identity across time was unambiguous, eliminating the need for alignment and facilitating annotation and quantification even as changes in peak position affected overlap with other peaks. Changes in peak position for organic acids in our samples were compared with reported titration curves^13,27,28^, in-house titrations for citrate (**Supplementary Fig. 4**), and Bruker AssureNMR software (Bruker Biospin, USA; **Supplementary Table 2**) to estimate pH of the sample at each timepoint. Our data indicate that the pH of the aerobic cultures began at 6.2-6.4, then dropped to 5.2-5.4 coincidentally with glucose consumption. Furthermore, this acidification reversed after glucose depletion at 6-7h, and pH increased to 5.5-5.7 by the end of our experiments. In the anaerobic samples, the pH decreased from 6.2-6.3 to 5.7-5.9. Although we did not perform high-resolution titrations for glutamate, succinate, and fumarate, their reported shifts were consistent with the trends for citrate (**Supplementary Table 2**).

Maintenance of characteristic differences in pH is well-accepted between organelles, the cytoplasm, and the extracellular milieu^29-31^. Filamentous fungi including *N. crassa*^32^ secrete large amounts of organic acids such as citrate, fumarate, and succinate, to acidify their extracellular environment^31,33,34^, and the two latter acids are taken up by carbon-limited *N. crassa*, with maximal uptake occurring around pH 5.5 ^31^.

### Activation of central carbon metabolism in aerobic conditions

Under aerobic conditions, glucose and trehalose were consumed within the first 6h while ethanol and lactate were produced. Glucose fell below our limit of detection by ^~^6h, after which it was depleted as the concentrations of ethanol and lactate plateaued (**Figs. 3a-b, 5**). The inverse trends between sugars and fermentation products indicate that most carbon was shunted to fermentation. In fact, we observed that most ^13^C in labeled glucose was channeled to ethanol and succinate (**Supplementary Fig. 5**), consistent with known *N. crassa* biochemistry^35,36^. Low levels of trehalose, another energy metabolite in the aerobic samples are consistent with vegetative growth, while increasing levels in the anaerobic samples may reflect a developmental shift to production of conidia in response to stress^37^.

**Figure 5.**
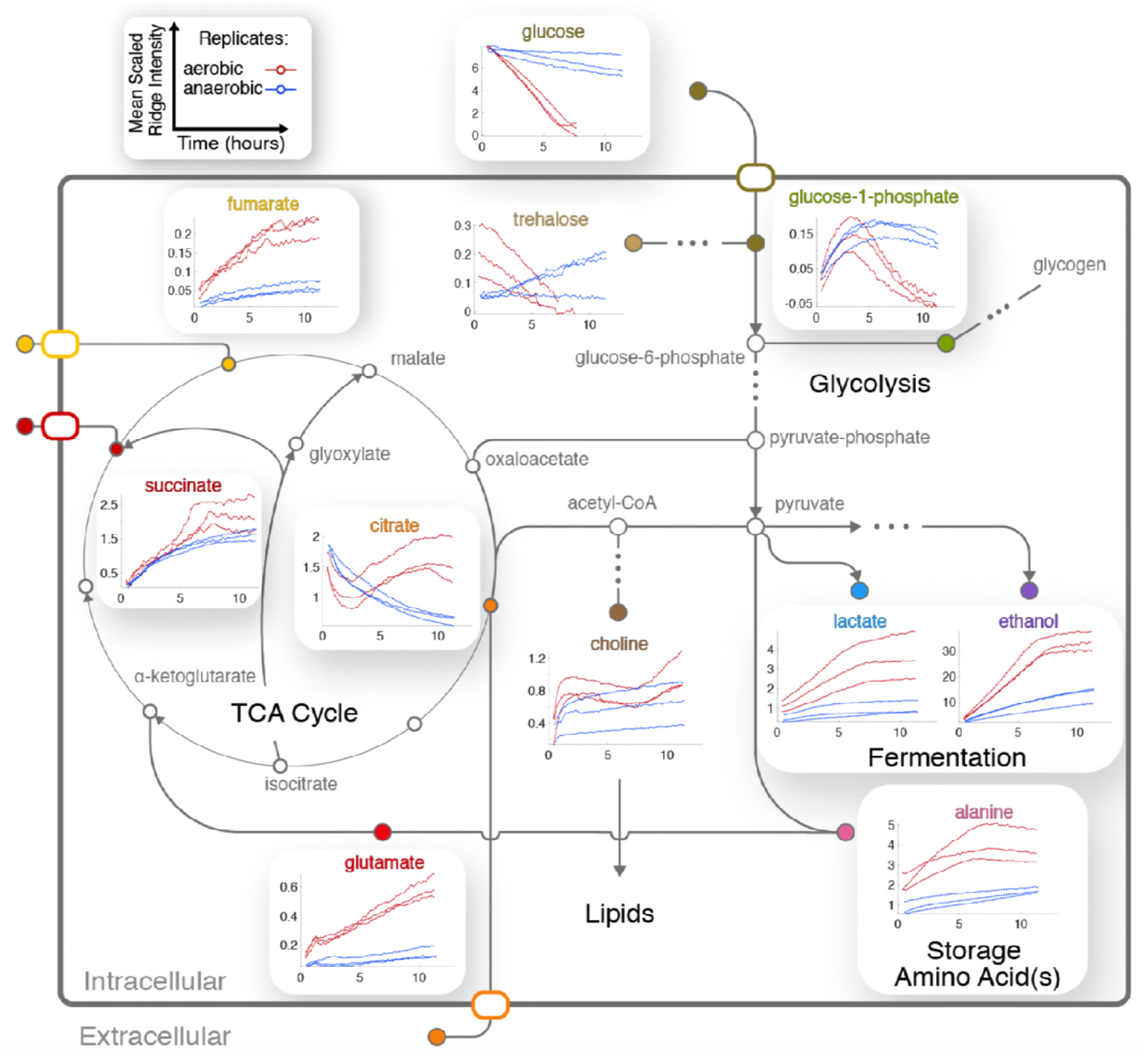
Integration of central metabolic pathways. Arrows correspond to one or more reactions, and nodes correspond to metabolites^39,40^. Nodes are filled for observed metabolites.

Four TCA cycle metabolites were detected in our experiments (**Figs. 3b-c**). Fumarate and succinate increased in the aerobic condition, and both accumulated slightly faster around 6h following glucose depletion and remained abundant (**Fig. 5**). This dynamic could be a result of glyoxylate cycle^38^ or mixed acid fermentation^39^ activation, and would not be observable without a continuous, densely sampled time series. Standard replicate averaging with extracted samples at different times would average out much of this detail. Next, one of the clearest signs of low oxygen levels in the anaerobic sample was the slight reduction in succinate compared to the drastic reduction in fumarate. Succinate levels in the aerobic condition are comparable to those in the anaerobic condition, while fumarate accumulates much more in the aerobic condition (**Fig. 5**). This is explained by the fact that conversion from succinate to fumarate depends on oxygen reduction in the electron transport chain^39,40^. Finally, citrate was abundant in the aerobic condition and followed a complex trend, while malate was observed in endpoint extracts (**Supplementary Fig. 6a**). Taken together, these trajectories constitute strong evidence for active glycolysis and fermentation in the presence of glucose and oxygen, while alternative pathways such as the glyoxylate cycle were active after glucose depletion. Similar trends with lower rates were observed in the anaerobic samples, except for differences in citrate and glucose-1-phosphate (**Fig. 5**).

### Interplay between amino acid, central carbon, and nitrogen metabolism

The dynamics of glutamate were different between the two conditions (**Fig. 5**). Glutamate accumulates while synthesis of glutamine is repressed in *N. crassa* in nitrogen-sufficient conditions^41^. We could not annotate glutamine with confidence because of overlap (**Supplementary Table 1b**). However, resonances consistent with glutamine increased after ^~^3h (**Supplementary Fig. 6b**), indicating potential nitrogen insufficiency in the aerobic culture. Arginine levels correspond to those of glutamate in the aerobic condition (**Supplementary Fig. 3**). Glutamate is produced from arginine degradation^38^; for instance, arginine has been reported as an abundant amino acid in extracted samples of actively growing *N. crassa* cultures^26,41^ and is thought to be catabolized to glutamate during conidiation^26^.

Trends for alanine (**Fig. 5**) and an unknown in the aliphatic region (**Fig. 4a**) were very similar to that of ethanol and lactate (**Fig. 5**), indicating that their metabolic fluxes are closely dependent on intermediates or energy produced by glycolysis and fermentation. This hypothesis is supported by the fact that alanine is synthesized from glutamate and pyruvate by alanine transaminase^41,42^. Glutamate levels increased and were unaffected by glucose, but alanine first accumulated and then decreased upon glucose depletion (**Fig. 5**). We conclude that alanine synthesis was limited by a lack of pyruvate caused by glucose depletion. Glutamate levels are maintained during starvation^38^, and Kanamori et al. (1982) suggested that alanine serves as a storage for pyruvate and nitrogen *via* glutamate in favorable conditions^41^. Therefore, the observed decrease in alanine suggests that it was utilized for pyruvate and glutamate when glucose concentrations were low in the aerobic condition (**Fig. 5**). Our CIVM-NMR data therefore supports glutamate as a hub between central carbon and amino acid pathways and confirms the maintenance of glutamate stores even under starvation.

### Complex trends reveal dynamics between energy storage and cell wall synthesis pathways

CIVM-NMR data revealed significant changes that preceded glucose depletion at ^~^6 h for compounds such as citrate, choline, adenosine, and valine, which all had similar trends in the aerobic condition (**Fig. 5**). Citrate decreased at the start of all experiments. Under aerobic conditions it began to accumulate again around 2.5 h and surpassed initial levels, while in anaerobic conditions it decreased at an exponential rate to a very low amount (**Fig. 5**). In the anaerobic samples, citrate was utilized while succinate accumulated to levels similar to those in the aerobic samples. However, fumarate levels remained low. Lack of oxygen could explain low rates of conversion from succinate to fumarate, while glyoxylate cycle activity can occur in anaerobic conditions^43,44^ and yields succinate and malate without fumarate.

*N. crassa* does not survive on citrate as a sole carbon source^45^, and to our knowledge extracellular citrate utilization has not been reported for *N. crassa*. However, citrate levels were observed well below the initial amount present in the media alone (9.74 mM) in both conditions, strongly indicating that external citrate was consumed in both experiments. Isotopic labeling experiments will more directly test this hypothesis.

Pyruvate and acetyl-CoA both serve as crossroads between major energy metabolites and lipids. Although we did not observe pyruvate and acetyl-CoA directly, most accumulating metabolites in pathways emanating from pyruvate exhibited strikingly similar trends (**Fig. 5**), suggesting flux through pyruvate. Curiously, citrate and choline did not follow this pattern, indicating activity from pathways that consume and replenish their pools. However, the rates of change of these metabolites were clearly opposed in both aerobic and anaerobic samples. This opposition suggests that flux from acetyl-CoA was being channeled differentially between citrate and choline synthesis and demonstrates a major carbon and energy exchange between central metabolism and lipid precursors^46^.

The accumulation of citrate and choline after glucose depletion in the aerobic samples was puzzling. Prior work has indicated that under low oxygen or glucose depletion *N. crassa* cells become vacuolated^47,48^. The synthesis of membranes for the vacuoles and their membranes under anaerobic conditions would explain the rise in choline. A concordant decrease in G1P at ^~^3h may indicate a shift of carbon flux to glycolysis from glycogen, caused by sensing of extracellular glucose levels^49^ or limitations of glycogen capacity. Glucose conversion to G6P is the first step of glycolysis^38^, which was clearly active in the first stages of our aerobic condition (**Fig. 5**). High levels of G6P drives its conversion by phosphoglucomutase to G1P^38^, which is converted by UDP-glucose pyrophosphorylase and UTP hydrolysis to the direct glycogen precursor UDP-glucose^38^. The latter is the rate-limiting step in glycogen synthesis, which is an endergonic process. If G6P levels were high and flux were shunted to glycogen, high levels of G1P would be expected. In fact, G1P levels increased in the aerobic samples until around 3 h then decreased, while UDP-Glucose was also observed. Therefore, we conclude that glycogen synthesis occurred in the first half of the aerobic experiments, but slowed or reversed after 3 h. This conclusion is further supported by the fact that glycogen synthesis occurs during high rates of growth in *N. crassa*, and wanes during slow growth^50-52^.

On the other hand, high levels of G1P are unlikely to be produced by the relatively low levels of glycolysis observed in the anaerobic samples and can indicate glycogen degradation. G1P accumulated to comparable levels in both conditions, but it remained in the anaerobic samples. Glycogen degradation is exergonic and releases G1P and glucose directly in an approximate 9:1 ratio^38^. Furthermore, UDP-Glucose was not observed in the anaerobic samples. Thus, high levels of G1P in the anaerobic conditions may indicate glycogen degradation.

Plots show the means of scaled peak/ridge intensities for a given compound in a given replicate over traceable times, where red and blue trajectories represent aerobic and anaerobic conditions, respectively. Arrows indicate typical reaction directions. The glyoxylate cycle is shown as a shunt through glyoxylate embedded in the TCA cycle.

The primary chitin cell wall building block UDP-N-acetylglucosamine (UDP-GlcNAc)^53^ increased in only the aerobic cultures (**Fig. 3, Supplementary Fig. 6c**), although overlap and low intensity prevented quantification. UDP-GlcNAc is synthesized via the unidirectional Leloir pathway^53^, and the only known uses for UDP-GlcNAc in *N. crassa* are chitin/cell wall biosynthesis and UDP-GalNAc production^53,54^. Filamentous fungi such as *N. crassa* produce chitinases^55^ and could utilize these for autolysis under stress conditions. However, if an increase in UDP-GlcNAc indicated cell wall degradation (i.e. due to stress or autolysis), those resonances would be expected to increase in the anaerobic condition; however, they were barely detected (**Fig. 3, Supplementary Fig. 6c**). Curiously, a recent study suggested that *N. crassa* utilizes alternative chitin catabolism pathways that would not result in increased GlcNAc-derived UDP-GlcNAc^56^. Considering the above dynamics, we conclude that resources were allocated between energy storage and cell wall synthesis pathways in glucose-rich conditions.

## DISCUSSION

We have demonstrated the use of CIVM-NMR to monitor metabolic dynamics in cells and whole microorganisms. An uninterrupted, high-resolution time series of NMR data allows observation of rapid but reproducible metabolic events. In contrast, using traditional studies of different replicates for each time point, the biological and technical variation often obscure details of dynamics. The lack of extraction removes a major source of technical variation found in typical MS and NMR metabolomics workflows and can facilitate inter-study comparisons.

NMR has relatively low sensitivity, but it is a quantitative and reproducible technique, and conventional NMR cryoprobes allow routine proton detection of compounds at concentrations as low as about 5 µM ^1^H. HR-MAS probes are generally less sensitive. However, the temporal dimension of CIVM-NMR data allows for more confident assignment of peaks with surprisingly low signal-to-noise ratios. By taking advantage of this unique property of CIVM-NMR data, we detected peaks as low as ^~^24-62 µM ^1^H (**Supplementary Fig. 7**). The sensitivity of CIVM-NMR is therefore particularly well-suited for observation of the major sources, sinks, and bottlenecks of metabolism in an organism or cells. For instance, an absolute quantification of 103 metabolites in *E. coli* revealed intracellular concentrations ranging from 0.13 µM to 96 mM^57^. As such, CIVM-NMR presents a unique opportunity to identify and monitor time-dependent metabolic resources and products in metabolic engineering of microbial organisms.

Only 20-70 µL of sample is needed with no sample preparation to yield an entire time series for various metabolites, and the sample can be used in downstream *in vivo* or chemical analyses following NMR data collection. These factors make CIVM-NMR ideal for scarce samples that would not otherwise be possible to study by time-series metabolomics^3^. With an internal rotor radius of 1.4 mm spinning at 6000 Hz, our samples experienced up to 200,000 x g of acceleration. As sedimentation was not observed, it is possible that a low relative density of *N. crassa* mycelia compared to the media may have resulted in a lower effective radius of rotation. While some samples, including the leukemia cells in **Figure 1**, are less stable at high spinning rates, microorganisms such as *E. coli* and *S. cerevisiae* can grow under different amounts of hypergravity, even with cellular and organellar sedimentation^58^. Furthermore, methods have been developed to obtain HR-MAS data with slow spinning^59^, which could allow monitoring in ^~^1500 x g or less. The lack of perfusion and a limited sample volume are both factors that need to be considered with regard to nutrient depletion and waste accumulation. Lastly, identification of spectral features and deconvolution of overlap are still challenging in any NMR or LC-MS metabolomics study. However, temporal continuity clearly provides information (e.g. as seen in **Fig. 3** and **Supplementary Fig. 7**) that will be helpful in addressing these problems.

Full utilization of the time series data from CIVM-NMR will require a modeling-based method, and our data underscore the need for accurate and experimentally-based kinetic models of metabolism. With the potential for <4-minute resolution by using fewer scans before saving fids at the cost of signal-to-noise ratio, CIVM-NMR provides a unique opportunity for probing flux changes as well as allosteric regulation^60^ with kinetic models^1,11^ for abundant metabolites. Each replicate can be seen as a single, complete model with different initial conditions, which is significantly better than a time series of averages. Previous real-time methods^11,13^ have equal or greater temporal resolution at the expense of disadvantages such as being destructive^1^, limitation to cell suspensions^1,13^, primarily measuring the media^8,13^, or having combined biological and technical variance. CIVM-NMR minimizes noise by eliminating sampling and extraction variance. Batch effects for each replicate are eliminated since all experimental and NMR parameters are consistent across timepoints. Analytical drift is eliminated because the detector never contacts the samples, and the sample is not perturbed by measurement. These factors in turn facilitate optimization of modeling parameters^61^.

CIVM-NMR can also allow continuous monitoring of the metabolic state before, during, and after a range of environmental and genetic perturbations. For instance, we observed a shift from a glucose-rich environment to starvation at ^~^6h. Sequential utilization of alternative carbon sources such as quinic acid^62^ is a natural extension of this work. In studies involving targeted pathways in mammalian cells, CIVM-NMR can facilitate the testing of mutants, gene knockdowns, or small molecule substrates, inhibitors, or activators. We are exploring the use of alternative gas mixtures for spinning the rotor, allowing adjustment of O^2^:CO^2^:N^2^ ratios during experiments. This presents another environmental shift and facilitates monitoring of cells (e.g. mammalian) which require controlled gas compositions. Real-time temperature control could also be used to probe temperature shifts or assess the effects of a temperature-sensitive mutation on the metabolism of an organism. Perturbations such as these will be critical to exploring and refining dynamics in kinetic models with an empirical basis.

## METHODS

Methods and any associated references are available in the online version of the paper. Any Supplementary Information and Source Data files are available in the online version of the paper. All code and data are freely available at https://github.com/artedison/Edison_Lab_Shared_Metabolomics_UGA and http://www.metabolomicsworkbench.org/.

## ACKNOWLEDGMENTS

We thank the A. Simpson lab for discussions and advice on drilling holes in rotor caps, M. Case, Clifford Slayman, and Z. Lewis for discussions on *Neurospora* stress and metabolism, and J. Walejko and G. Gouveia for discussions on method development. L. Morris provided guidance in bash scripting for automation of NMRPipe commands. B. Schuttler and L. Mao provided stimulating discussions on *N. crassa* modeling.

This work was primarily supported by NSF 1713746. ASE was also supported by the NSF ERC 1648035 (CMaT) and the Georgia Research Alliance.

## AUTHOR CONTRIBUTIONS

M.T.J. and Y.W. contributed equally to this work. M.T.J., A.S.E., and J.A. designed the project. M.T.J., Y.W., and A.S.E. wrote the manuscript. M.T.J and Y.W. wrote the scripts and analyzed the results. M.T.J. prepared *N. crassa* samples and extracts. M.T.J., A.S.E., and J.G. performed *in vivo* metabolite measurements and processed the data. F.T. collected and annotated 1D and 2D experiments on extracts. T.I., A.H., and J.G. designed and performed *in vivo* carbon-labeled measurements on human cells.

## Materials and Methods

### Human Leukemia Cell Culture and Preparation for HR-MAS NMR

The human BC-CML cell line K562 was obtained from ATCC, and cell line authentication testing was performed by ATCC-standardized STR analysis to verify their identity. After cell counting and washing with PBS, K562 cells were resuspended and labeled in a custom-made Iscove’s modified Dulbecco’s Medium (IMDM) without BCAAs supplemented with 10% dialyzed FBS, 100 IU/ml penicillin, 100 μg/ml streptomycin and the following amino and keto acids. For ^13^C-KIV tracer experiment, isoleucine, leucine and valine were supplemented at 170 µM. For ^13^C-valine tracer experiment, isoleucine, leucine and KIV were added at 170 µM. Cell suspension (54 µl) was loaded in a clean 4 mm diameter zirconia HR-MAS rotor (Bruker BioSpin), and then either [(U)-^13^C]-ketoisovalerate or [(U)-^13^C]-valine solution in D_2_O was added to a final concentration of 170 µM. Rotor was closed with a Kel-F rotor cap (Bruker BioSpin).

### Preparation of Growth Media and Slants for *N. crassa*

Ingredients for Vogel’s Media (3 % Glucose)

(Glucose, 0.334 M; Biotin, 0.614 µM; Arginine, 1.95 mM; Na_3_ Citrate, 9.74 mM; KH_2_PO_4_, 36.7 mM; NH_4_NO_3_, 25.0 mM; MgSO_4_, 0.811 mM; CaCl_2_, 0.680 mM; ZnSO_4_, 34.8 µM; Fe(NH_4_)_2_(SO_4_)_2_, 5.10 µM; CuSO_4_, 2.00 µM; MnSO_4_, 0.592 µM; H_3_BO_3_, 1.62 µM; Na_2_MoO_4_, 0.413 µM) were dissolved in ddH_2_O in a large glass bottle, filter-sterilized (0.22µm Steritop Threaded Bottle Top Filter, 500mL, Millipore EMD), stirred using a magnetic stir bar, then aliquoted into clean, sterile 500-mL bottles. Ingredients for Vogel’s Media with Agar (same as above, with the addition of 1.5 % agar, w/v, and using 1.5 % Glucose, w/v) were combined in a beaker. Agar was dissolved by heating in a microwave oven. The dissolved mixture was aliquoted to 15-mL or 5-mL glass test tubes, stoppered with cotton, and sterilized by autoclaving.

### Vogel’s Media for NMR and Wash Solution

2X Vogel’s Media (minus glucose), DSS solution, and D_2_O were combined to make a concentrate, which was split into two aliquots.

To prepare Vogel’s Media for NMR (1.5 % Glucose), filter-sterilized D-glucose solution (0.5mg/µL) was added to the smaller aliquot to a final composition of Glucose, 0.167 M; DSS, 1mM; Biotin, 0.614 µM; L-arginine, 1.95 mM; Na_3_ Citrate, 9.74 mM; KH_2_PO_4_, 36.7 mM; NH_4_NO_3_, 25.0 mM; MgSO_4_, 0.811 mM; CaCl_2_, 0.680 mM; ZnSO_4_, 34.8 µM; Fe(NH_4_)_2_(SO_4_)_2_, 5.10 µM; CuSO_4_, 2.00 µM; MnSO_4_, 0.592 µM; H_3_BO_3_, 1.62 µM; Na_2_MoO_4_, 0.413 µM in 95 ddH_2_O/5 D_2_O (v/v). Wash Solution was prepared by adding ddH2O in place of D-glucose solution to the larger aliquot.

### Preparation and Storage of *N. crassa* Conidial Suspension

A frozen bd1858(A) stock obtained by the Fungal Genetics Stock Center^1^ was used to inoculate two Growth Slants (Vogel’s Media Agar, 1.6% Glucose w/v, 3 mL in 15 mL glass test tubes stoppered with sterile cotton plugs). These were incubated for 2 days at 30°C, then placed under a benchtop lamp at 25 °C for 2 days to induce maturation of conidia. Conidia were collected from both tubes sequentially by suspension in 12 mL Vogel’s Media (no glucose) and filtration through sterile cotton. Concentration of the resulting conidial suspension was found to be 6.47 × 10^7^ cells/mL using a Nexus Cellometer Auto 2000 (Nexelcom Bioscience; Lawrence, MA, USA). The conidial suspension was kept at 4 °C over the course of the experiments (4 weeks).

### Growth of *N. crassa* Mycelia

Vogel’s Media (50 mL, 3 % Glucose w/v) in a 250-mL Erlenmeyer flask, covered in aluminum foil was inoculated under aseptic conditions with Conidial Suspension to a total concentration of 2.7 × 10^4^ cells/mL, (21 µL Conidial Suspension). Liquid cultures were grown with orbital shaking (^~^237 rpm) at room temperature (^~^25 °C) under constant cool white light (7 µmol L^−1^ s^−1^ m^−2^) for 32h. At that point mycelia consistently formed a single, cohesive mass. The entire culture was transferred to a 50 mL Conical Tube (Sarstedt; Newton, NC, USA) for transport to the NMR facility (15-30min).

### Preparation of *N. crassa* Mycelia

Under aseptic conditions, a section of mycelium from the edge of the main mycelial mat was cut off using a sterile tube cap and trimmed to fit the volume of approximately 126 µL using a pre-marked microcentrifuge tube. Mycelia were handled from this point using clean, sterile tweezers (cleaned with 70 % EtOH on a lint-free single-ply lab tissue (kimwipe) and dried in an aseptic environment).

The section of mycelium was then patted dry on autoclaved filter paper (Whatman Filter Paper #3; GE Healthcare, USA) atop a layer of folded kimwipes, and was washed by placing in a sterile microcentrifuge tube containing 1 mL Wash Solution and vortexing briefly (^~^10 s) until the mycelium had fully absorbed the media. Washing was repeated with fresh Wash Solution for a total of 4 washes. The mycelium was reduced to ^~^63 µL (0.9 × volume of rotor + plug), measured in a second microcentrifuge tube pre-marked to that volume. The mycelium was pat-dried in a sandwich of sterile filter paper folded into kimwipes, pressing firmly three times (until no liquid spots were visible on the filter paper). The dried mycelium was then weighed in a separate microcentrifuge tube. The dry mycelium was 9.04-10.13 mg in our experiments (µ = 9.62 mg; SD = 0.32 mg). We observed a reduction in mass of ^~^30 % as conidia, loose filaments, and other debris are removed along with waste products and glucose during wash steps. In our hands, the prep process took between 4 and 13 min., during which time the organism was immersed in a low-glucose environment.

### Loading *N. crassa* Mycelia into the Rotor

The dried, weighed mycelium was then placed in a microcentrifuge tube containing fresh Vogel’s Media for NMR (500 µL, 1.5 % Glucose), and vortexed briefly until the mycelium had fully absorbed the media. The mycelium was then transferred to a third, pre-marked microcentrifuge tube (63 µL). By adding/removing media, the volume was adjusted to the 63 µL volume mark. Sterile tweezers were used to transfer the mycelium to a clean 4 mm diameter zirconia rotor (Bruker BioSpin) cleaned by rinsing with bleach solution, tap water, 70 % ethanol, tap water, and ddH_2_O × 4). The mycelium was pushed to the bottom, taking caution not to lose liquid. The remaining liquid in the tube was added to the rotor and one tweezer prong was used to position the mycelium to remove larger air bubbles, although small bubbles occurred with no issues in the NMR. A Teflon sealing plug (Bruker BioSpin) was then inserted to ^~^2 mm below the edge of the rotor. For the aerobic condition, a Kel-F rotor cap (Bruker BioSpin) modified with a 0.016-inch diameter hole drilled using a lathe was lined on the inside with three layers of Rayon breathable microplate sealing tape (QuickSeal Breathable Film, Thomas Scientific, USA) to prevent spore escape. The cap was fully inserted to push the sealing plug into its final position. The cap was then removed, and the insides of the cap and plug were inspected to ensure that no liquid was lost and that an airspace existed between the plug and the sample. The rotor was then re-capped, the bottom edge marked with a permanent marker, and dropped into the bore of the magnet (cap facing up). In our hands, this process typically takes 15-30 min. For the anaerobic condition, media was added to fill all airspaces and an unmodified cap was used to prevent gas exchange. For the ^13^C labeling experiment in partially anaerobic conditions, an airspace was left and fresh Vogel’s Media for NMR was prepared with 3% (w/v) ^13^C-labeled Glucose (99% labeled; Cambridge Isotope Laboratories; Tewksbury, MA, USA) was used in place of 1.5% (w/v) glucose.

### NMR Parameters

For human ML cell experiments, a hsqcetgpsisp.2 gradient HSQC run as a 1D experiment was used with the following parameters: Data Points: 7272. Dummy Scans: 4 at the beginning of the run. Number of Scans: 128 /timepoint. O1 offset: 4.699 ppm. O2 offset: 30.000 ppm. Acquisition Time 0.3999600 s. Delay: 1.5 s. fid Resolution: 2.500250 Hz. Receiver Gain: auto (101). Temperature: 298 K = 25 °C. Spinning Speed: 3100 Hz.

A standard noesypr1d protocol (Bruker) was used for *N. crassa* non-labeled real-time metabolomics measurements. The following parameters applied to all samples and timepoints: Data Points: 42856. Dummy Scans: 8 at the beginning of the run. Number of Scans: 64 /timepoint. Spectral Width 11904.762 Hz. Acquisition Time 1.7999520 s. Delay: 1.5 s. fid Resolution 0.555570 Hz. Receiver Gain: auto (101). Temperature: 298 K = 25 °C (calibrated using a deuterated methanol standard^2^). The following parameters were optimized for each sample: O1 offset for water suppression: 2817.24 - 2818.24 Hz. PWL9 water suppression power: 43.87 - 44.42 dB (µ = 44.23 dB, SD = 0.19 dB). P1 pulse width: 12.49 - 13.30 µs (µ = 12.78 µs, SD = 0.29 µs). Spinning Speed: 6000 Hz. *Notably, this variation in pulse width between samples manifested as a difference in temporal resolution (i.e. longer pulse widths resulted in time points slightly farther apart). The effect was measurable (on the order of minutes) over hundreds of measurements. The average experiment took 4.23min +/− 0.004min (SD)*.

For measurement of ^13^C in the labeled glucose experiment, a modified hmqc with additional phase cycling was used. The following parameters were used: Data Points: 3636. Dummy Scans: 8 at the beginning of the run. Number of Scans: 64 /timepoint. O1 offset: 2826.24 Hz. O2 offset: 12070.62 Hz. Acquisition Time 0.2545200 s. Delay: 1.5 s. fid Resolution 3.928964 Hz. Receiver Gain: auto (101). Temperature: 298 K = 25 °C. Spinning speed: 6000 Hz.

All Bruker parameter files are available with the raw data at http://www.metabolomicsworkbench.org/.

### Automated Data Acquisition and Post-experiment Sample Preparation

For human ML cells, spectra were collected sequentially using the multizg command in TopSpin (v4.0.1; Bruker).

For *N. crassa* samples, the noesypr1d experiment, optimized for the sample, was imported into IconNMR in TopSpin (v4.0.1; Bruker). The solvent was set to “D2O_H2O+salt”. The “iterate” command was used to queue 1024 identical, sequential noesypr1d experiments (each taking ^~^4.6 min) on a Bruker NEO equipped with a 4-mm CMP probe. Dummy scans were only implemented for the first experiment. Experiments generally ended after ^~^12 h, though some were allowed to continue as long as 37 h. By spinning *N. crassa* at 6 KHz, spinning sidebands^3^ were eliminated in the spectral region of 0-10 ppm. At the end of each run, the mycelia were transferred from the rotor to a sterile microcentrifuge tube with clean, sterile tweezers. All liquid from the rotor was also transferred to the tube. This was either extracted and assessed for growth immediately, or was allowed to sit on the bench for one day.

### Survival Assessment

Sterile tweezers were used to tear a piece of mycelium from the rotor contents; this was used to inoculate a growth slant. All growth slants were assessed for 24 h or longer post-inoculation for growth. Photographs were taken using a 16MP digital camera on an LG G5 cell phone.

### Extraction

The remaining rotor contents were transferred with a pipette to a microcentrifuge tube containing a mixture of zirconia beads (1 mm, 167 µL or ^~^375 mg; 0.7 mm, 334 µL or ^~^1314 mg; 500 µL total) on dry ice. The old tube was rinsed by briefly vortexing with 800 µL MeOH (80 % in ddH_2_O), which was added to the beads. This mixture was either processed immediately or frozen on dry ice for up to 3 days. Contents were twice homogenized on dry ice for 180 s @1800 rpm using a MP FastPrep 96 (MP Biomedical; USA) adapted for microcentrifuge tubes, adding dry ice each time. The homogenate was centrifuged at 14k rpm at 4 °C for 5 min (18220 x g; Centrifuge 5417C; Eppendorf, USA). The supernatant was transferred to a separate microcentrifuge tube and kept on dry ice while the pellets were back-extracted with 500 µL MeOH (80 %), homogenized once for 180 s @1800 rpm, and centrifuged an additional 5 min. Supernatants from both extractions were combined, then dried to completion in a CentriVap Concentrator/CentriVap Cold Trap −105 °C system (Labconco, Kanasas City, MO, USA) for 4-6 h. Pellets for two samples were combined during resuspension in D_2_O (DSS, 1/9 mM) for each condition. Two replicates from each condition were pooled and pipetted into 1.5 mm NMR tubes (Norell; Morganton, NC, USA).

### Annotation

For each pooled sample representing the anaerobic and the aerobic conditions, noesypr1d, ^13^C-HSQC, TOCSY, and ^13^C-HSQC-TOCSY spectra were collected on a 600 MHz Bruker magnet equipped with a 5mm cryoprobe and an Avance III HD console at the University of Georgia NMR Facility. 2D data were processed in nmrPipe (System Version 9.4 Rev 2017.340.17.07 64-bit) and submitted to COLMARm^4^ for putative compound identification. After manual inspection, metabolites were assigned a confidence level ranging from 1 to 5, with 5 being the highest. The scale is defined^5^ as follows: (1) putatively characterized compound classes or annotated compounds, (2) matched to literature and/or 1D reference data such as HMDB^6^ and BMRB^7^ (3) matched to HSQC, (4) matched to HSQC and validated by HSQC–TOCSY (COLMARm^4^), and (5) validated by spiking the authentic compound into sample. Identifications from extracted 1d spectra were manually mapped to real-time *in vivo* noesypr1d data. An additional score was assigned to each mapped compound: 0 (unannotated), 1 (annotated only), 2 (qualitatively assessed), or 3 (relatively quantifiable) in the real-time data. This score depended on number of observed peaks, baseline, peak overlap, and sensitivity. Both metabolite confidence levels are reported in Table S1. All raw and processed data files are available at http://www.metabolomicsworkbench.org/ and matching can be run on COLMARm^4^ directly.

### Batch Processing in nmrPipe for *in vivo* NMR Data

Parameters were optimized based on agreement between spectra from several time points for a given sample. A custom bash script ran nmrPipe^8^ using the optimized parameters on all spectra for a given sample. This script included all necessary nmrPipe commands for file conversions and NMR data processing. In brief, the following were implemented: line broadening, Fast Fourier Transform, 0- and 1^st^-order phasing, end removal, and baseline correction using automatic polynomial fitting. All raw data, parameter files and code are available at http://www.metabolomicsworkbench.org/.

### Additional Processing in MATLAB for *in vivo* NMR Data

For each sample, custom scripts were written in MATLAB R2017b (The MathWorks, Inc., Natick, Massachusetts, USA), to load the processed spectra, ppm vectors, and measurement start times from .ft and .acqus files. Spectra were then referenced to DSS semi-automatically, stored as a matrix, and saved as a MATLAB workspace in .mat format.

Using custom MATLAB scripts, .mat files from individual experiments were combined into a “sampleData” structure. Metadata (e.g. condition, pulse width, time shift between inoculation and start time) were added to each sample by manual entry or by automated retrieval from the Bruker acqus files for each sample. Spectral ends outside of [-0.5,10] ppm were removed. The spectral region containing the water signal [4.7,5] ppm was replaced by zeros. Measurements for time points >11 h were removed in all experiments for consistency. Each spectrum was normalized to its DSS peak intensity as a formal step to allow for relative quantification. Finally, every three spectra were summed starting from the first timepoint for improved signal-to-noise. The resulting structure was saved as a .mat file (^~^2 Gb). All data and scripts are available at http://www.metabolomicsworkbench.org/ and at https://github.com/artedison/Edison_Lab_Shared_Metabolomics_UGA.

### Relative Quantification of NMR Resonances

A combination of a Gaussian smoothing filter with user-defined sigma in the ppm and time dimensions and peak picking script was used to identify peak maxima for a given region of ^~^0.5-1 ppm in a given sample, allowing some noise to be picked. Agglomerative clustering based on single linkage of Euclidean distances was then used to cluster the picked points in the chemical shift (ppm), time, and intensity space. Weights for each dimension in the clustering, as well as the number of clusters, were manually optimized for each region and sample. Clusters were quality-controlled by interactive visual inspection.

If multiple ridge points existed for the same time, the one with highest intensity was retained. Peak positions at temporal gaps were estimated using linear interpolation between the two closest existing ridge points. Ridges on the smoothed data were mapped to the unsmoothed data for each time point by choosing the maximum within a small window around the peak position obtained from the smoothed data.

A window size of 10 indices (^~^2.9 × 10^−3^ ppm) worked for all but a few ridges, whose optimal mapping windows ranged between 6 and 60 indices (between 1.7 × 10^−3^ and 1.7 × 10^−2^ ppm). All ridges were visually inspected for good tracing, well-defined peaks, and minimal overlap by plotting on real spectra.

To combine the trend information from multiple ridges annotated to the same compound, intensities of constituent ridges were scaled such that the ridge means across the time points shared by the highest number of ridges were equal. Lastly, the mean across scaled ridges at each time point was taken, yielding a single composite trajectory for each compound.

### Titration of a Citrate Standard for Estimation of *in-vivo* pH Changes

A 10 mM solution of citric acid (A104-500; Fisher Scientific, USA) containing 1mM DSS reference standard was prepared, and 600 µL were added to a 5 mm NMR tube (Norell; Morganton, NC, USA). The pH of the solution was adjusted in-tube in ^~^0.25 pH increments by addition of 0.5-2 µL volumes of dilutions of concentrated NaOH and HCl and four rounds of inversion and vortex mixing. For each pH point, exact pH was measured in-tube using a calibrated accumet AB150 pH meter (Fisher Scientific, USA), then a 1D noesypr1d spectrum was collected (DS = 2; NS = 16) on a 600 MHz Bruker magnet equipped with a 5mm cryoprobe and an Avance III HD console at the University of Georgia NMR Facility. Data were phased and referenced to DSS and in TopSpin (v3.5pl7; Bruker). Custom Matlab scripts were used to obtain the most upfield citrate peak position for each pH. A 3^rd^-order polynomial was fit to the positions (R^2^ > 0.99) and used with the ridge belonging to the same peak to estimate the pH of each culture at each timepoint.

